# Metabolic Adaptability and Nutrient Scavenging in Toxoplasma gondii: Insights from Ingestion Pathway-Deficient Mutants

**DOI:** 10.1101/2024.11.27.625683

**Authors:** Patrick A. Rimple, Einar B. Olafsson, Benedikt M. Markus, Fengrong Wang, Leonardo Augusto, Sebastian Lourido, Vern B. Carruthers

**Affiliations:** Department of Microbiology and Immunology, University of Michigan Medical School, Ann Arbor, MI, USA; Whitehead Institute for Biomedical Research, Cambridge, MA, USA; Department of Pathology, and Microbiology, and Immunology, University of Nebraska Medical Center, Omaha, NE, USA

## Abstract

The obligate intracellular parasite *Toxoplasma gondii* replicates within a specialized compartment called the parasitophorous vacuole (PV). Recent work showed that despite living within a PV, *Toxoplasma* endocytoses proteins from the cytosol of infected host cells via a so-called ingestion pathway. The ingestion pathway is initiated by dense granule protein GRA14, which binds host ESCRT machinery to bud vesicles into the lumen of the PV. The protein-containing vesicles are internalized by the parasite and trafficked to the Plant Vacuole-like compartment (PLVAC), where cathepsin protease L (CPL) degrades the cargo and the chloroquine resistance transporter (CRT) exports the resulting peptides and amino acids to the parasite cytosol. However, although the ingestion pathway was proposed to be a conduit for nutrients, there is limited evidence for this hypothesis. We reasoned that if *Toxoplasma* uses the ingestion pathway to acquire nutrients, then parasites lacking GRA14, CPL, or CRT should rely more on biosynthetic pathways or alternative scavenging pathways. To explore this, we conducted a genome-wide CRISPR screen in wild-type (WT) parasites and Δ*gra14*, Δ*cpl*, and Δ*crt* mutants to identify genes that become more fitness conferring in ingestion-deficient parasites. Our screen revealed a significant overlap of genes that become more fitness conferring in the ingestion mutants compared to WT. Pathway analysis indicated that Δ*cpl* and Δ*crt* mutants relied more on pyrimidine biosynthesis, fatty acid biosynthesis, TCA cycle, and lysine degradation. Bulk metabolomic analysis showed reduced levels of glycolytic intermediates and amino acids in the ingestion mutants compared to WT, highlighting the pathway’s potential role in host resource scavenging. Interestingly, ingestion mutants showed an exacerbated growth defect when grown in amino acid-depleted media, suggesting a role for the *Toxoplasma* ingestion pathway during nutrient scarcity.

**Importance:** *Toxoplasma gondii* is an obligate intracellular pathogen that infects virtually any nucleated cell in most warm-blooded animals. Infections are asymptomatic in most cases but people with weakened immunity can experience severe disease. For the parasite to replicate within the host, it must efficiently acquire essential nutrients, especially as it is unable to make several key metabolites. Understanding the mechanisms by which *Toxoplasma* scavenges nutrients from the host is crucial for identifying potential therapeutic targets. Our study highlights the function of the ingestion pathway in sustaining parasite metabolites and contributes to parasite replication under amino acid limiting conditions. This work advances our understanding of the metabolic adaptability of *Toxoplasma*.

## Introduction

The obligate intracellular parasite *Toxoplasma gondii* infects up to 30% of the world’s human population^1^. This success expands beyond humans as *Toxoplasma* can infect any nucleated cells in most warm-blooded animals. The extremely large host range suggest that *Toxoplasma* has extremely versatile mechanisms to support its growth.

Like other intracellular pathogens, *Toxoplasma* must derive its nutrients from the host cell it infects^2^. This is achieved through a combination of scavenging^3,4^ and host cell manipulation^5^ to complement de *novo* biosynthesis^6^. To scavenge material from the host cell *Toxoplasma* must bypass the parasitophorous vacuole membrane (PVM), which protects the parasite from intrinsic host defenses^7^. Scavenging of small soluble metabolites is facilitated by diffusion across the PVM through the nutrient pore proteins GRA17, GRA23, GRA47, and GRA72^8–11^. Once in the PV, transporters on the parasite plasma membrane mediate uptake into the parasite cytosol for utilization^12^. Lipids are scavenged by sequestering host-derived vesicles within the PV and transfer of lipid material to the parasite by unknown mechanisms^13–15^. Another potential nutrient acquisition route for *Toxoplasma* is the ingestion pathway, which entails parasite uptake of host cytosolic material. Using the secretory protein GRA14, *Toxoplasma* co-opts the host’s Endosomal Sorting Complex Required for Transport (ESCRT) machinery at the PVM to generate vesicles that the parasite internalizes, possibly at a specialized site of endocytosis, the micropore^16–18^. The ingested material traffics through the endosome-like compartment toward the PLVAC^19^. Once inside the PLVAC, the ingested proteins are degraded by TgCPL^20^, the major protease of the PLVAC^21,22^, and the digested products are likely liberated by a small peptide/amino acid transporter, the chloroquine resistance transporter (TgCRT)^23,24^ (Fig. 1A). Despite previous observations demonstrating that the ingestion pathway facilitates the acquisition of host cytosolic proteins, the exact reason for its employment by *Toxoplasma* remains unknown, particularly because disrupting this pathway has little impact on parasite fitness *in vitro*^23^.

**Figure 1.**
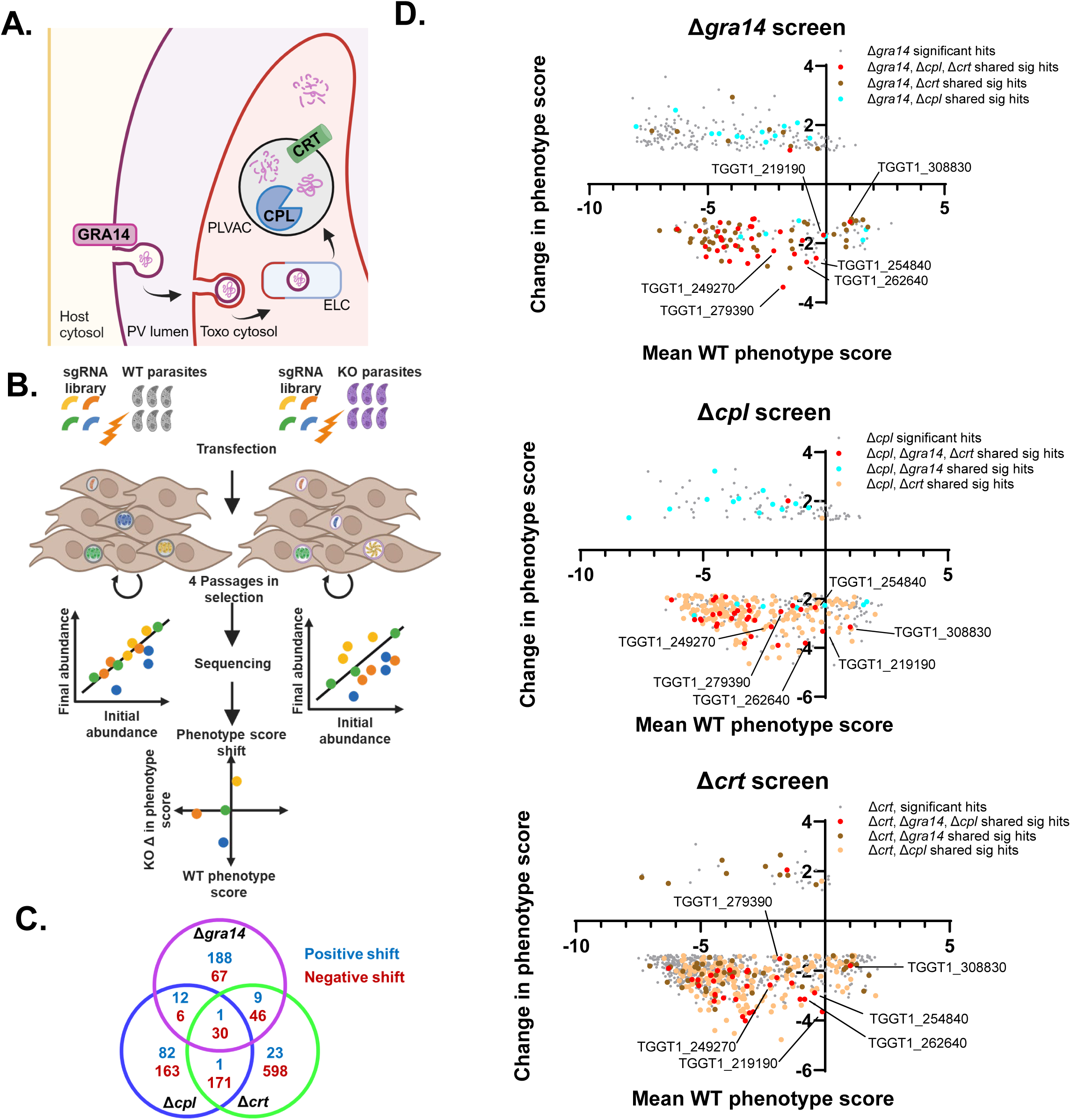
Genome-wide CRISPR screen identifies fitness conferring genes related to the ingestion pathway. **(A)** Schematic of the ingestion pathway wherein GRA14 is at the PVM to facilitate uptake of host cytosolic protein, the ingested material is trafficked through the parasite’s endosome like-compartment (ELC), delivered to the PLVAC for degradation by CPL and liberated by CRT. (**B)** Schematic of the CRISPR screens. 6×10^8^ parasites were transfected with a linearized plasmid library containing 10 sgRNA per gene of the *Toxoplasma* genome. Parasites were passed 4 times in pyrimethamine to select for integration of the sgRNA-expression cassettes subsequent disruption of the gene. sgRNA abundance was determined from the initial library and final population through Illumina sequencing and the phenotype score was calculated based on the top 5 sgRNAs for each gene. **(C)** Venn diagram indicating significant hits shared between samples. Cyan indicates genes shifted positively; red indicated genes shifted negatively. **(D)** Graphs depicting the significantly shifted hits for each screen, with the change in phenotype score on the y axis and the phenotype score for the WT screen on the x axis. Grey indicates hits pertaining to just the mutant on the graph, red indicates shared between all mutants, cyan indicates shared between Δ*gra14* and Δ*cpl*, tan indicates shared between Δ*cpl* and Δ*crt*, and gold indicates shared between Δ*gra14* and Δ*crt. n*=3 biological replicate screens for each strain.

We hypothesize that in the absence of the ingestion pathway, *Toxoplasma* compensates by tapping into alternative nutrient and metabolite sources such as through upregulation of biosynthesis pathways or increased utilization of other nutrient uptake systems. To test this, we employed a genome-wide CRISPR-mediated gene-disruption screen to identify genes that become more fitness conferring in ingestion-deficient mutants compared to WT parasites. By analyzing mutants that are deficient in three different steps of the pathway (Δ*gra14*, Δ*cpl*, and Δ*crt*), we aimed to uncover genes and pathways that compensate for the loss of ingestion, thereby shedding light on the broader metabolic adaptability and nutrient acquisition strategies of *Toxoplasma*.

## Results

Before performing a genome-wide CRISPR screen to identify genes that are differentially fitness conferring in ingestion mutants, we first generated ingestion mutant-strains in an RHCas9 strain background. We transfected RHCas9 parasites^25^ with sgRNAs targeting the first exon of *GRA14*, *CPL*, or *CRT* to generate loss-of-function INDEL mutants. We verified the disruption of *GRA14*, *CPL*, and *CRT* by Sanger sequencing across the sgRNA-targeted sites (data not shown) and western blotting (Fig. S1A–C). We also confirmed with an ingestion assay that RHCas9Δ*gra14* parasites have reduced uptake of host cytosolic protein (Fig. S1D).

A schematic of the screen is presented in Fig. 1B. Briefly, we screened the WT, Δ*gra14*, Δ*cpl*, and Δ*crt* strains using an established protocol^26^ with minor modifications, including passaging the library of mutant parasites four times instead of three times to give more time for fitness mutants to drop out of the population. After the final passage, genomic DNA was extracted and used to PCR amplify sgRNAs for sequencing and comparison to the sgRNAs in the initial library. We calculated the phenotype score for each gene in WT or mutant parasites by determining the Log2 fold change in relative abundance from the initial sgRNA pool to the final population. To minimize the effects of a bottleneck, we performed this analysis only for the top 5 most abundant sgRNAs per gene.

Despite a possible bottleneck (the final libraries were missing 40–60% of the sgRNAs), phenotype scores for the WT screen had an R² value of 0.85 compared to the original CRISPR screen in this strain^27^, which is comparable or higher than other published screens^28,29^ (Fig. S2A, B). We also observed a high correlation among WT replicates, suggesting reproducibility (Fig. S2C). Additionally, previously known essential and dispensable genes stratified along a phenotype score of -2.5, with essential genes having lower scores and dispensable genes having higher scores, consistent with previous screens^27^ (Fig. S2D).

After calculating the phenotype scores, we assessed for each gene whether the shift in phenotype score between WT and mutants was statistically significant (Table S1). We used a set of 497 genes expressed only in the sexual stages of the parasite as a reference cohort of dispensable genes to account for random variation in phenotype scores in WT and mutant parasites^30^. We applied two criteria for significance: first, identifying genes with a shift greater than 2 standard deviations from the mean shift among the sexual stage genes and second, performing a Wilcoxon-signed rank test using the shift in sexual stage genes as the control group to identify significantly shifted genes (FDR <0.01). Genes meeting both criteria were labeled as significantly shifted.

This analysis identified 149, 307, and 845 genes with significantly lower phenotype scores in Δ*gra14*, Δ*cpl*, and Δ*crt* mutants, respectively, and 210, 96, and 34 genes with significantly higher phenotype scores, respectively (Fig. 1C–D). We looked for hits that are shared among all three mutants and found 30 genes that become significantly more fitness conferring in all three mutants compared to WT (Table 1).

**Table 1.** Shared genes from Δ*gra14*, Δ*cpl*, and Δ*crt* fitness conferring hits.

| Gene ID | Product description | LoPIT | WT<br>mean<br>phen<br>score | $\Delta gra14$<br>mean<br>phen<br>score | $\Delta cpl$<br>mean<br>phen<br>score | $\Delta crt$<br>mean<br>phen<br>score | Sidik et<br>al 2016<br>phen<br>score |
| --- | --- | --- | --- | --- | --- | --- | --- |
| TGGT1_308830 | dual specificity protein phosphatase | N/A | 1.0107 | -0.2579 | -2.1373 | -0.784 | -0.17 |
| TGGT1_219190 | nuclear movement protein domain containing protein | cytosol | -0.1091 | -1.8295 | -3.4267 | -3.753 | -1.68 |
| TGGT1_254840 | tetratricopeptide repeat-containing protein | N/A | -0.4191 | -2.923 | -2.7681 | -3.305 | -1.4 |
| TGGT1_262640 | ApiCox23 | mitochondrion - membran | -0.8214 | -3.4588 | -4.6243 | -3.9658 | -3.49 |
| TGGT1_221320 | acetyl-CoA carboxylase ACC1 | apicoplast | -1.0052 | -2.9117 | -3.4354 | -4.1521 | -3.6 |
| TGGT1_286530 | ApiCox24 | mitochondrion - membran | -1.3416 | -3.702 | -3.6152 | -3.8367 | -2.82 |
| TGGT1_279390 | proliferation-associated protein 2G4, putative | nucleus - non-chromatin | -1.8223 | -5.3071 | -4.3315 | -3.3465 | -0.88 |
| TGGT1_288880 | hypothetical protein | N/A | -1.9415 | -3.5545 | -5.8399 | -4.2182 | -4.6 |
| TGGT1_249270 | protein disulfide isomerase-related protein (provisional), puta | apicoplast | -2.2093 | -4.4655 | -5.3494 | -4.781 | -4.56 |
| TGGT1_235398 | hypothetical protein | mitochondrion - soluble | -2.8858 | -5.3206 | -5.7581 | -6.5397 | -4.79 |
| TGGT1_267560 | folate-binding protein YgfZ protein | mitochondrion - soluble | -3.035 | -4.2071 | -6.5718 | -6.731 | -3.34 |
| TGGT1_254135 | hypothetical protein | N/A | -3.1081 | -4.2942 | -5.5791 | -5.0898 | -2.94 |
| TGGT1_220940 | ribosomal RNA methyltransferase (FtsJ) family protein | N/A | -3.1345 | -4.5725 | -5.8777 | -5.7592 | -2.38 |
| TGGT1_313140 | isocitrate dehydrogenase | mitochondrion - soluble | -3.1462 | -4.99 | -5.4087 | -5.1844 | -3.04 |
| TGGT1_254530 | hypothetical protein | N/A | -3.2036 | -4.6322 | -6.0362 | -7.2143 | -4.69 |
| TGGT1_273840 | brix domain-containing protein | nucleus - non-chromatin | -3.3115 | -5.9955 | -7.1283 | -7.1604 | -3.18 |
| TGGT1_319930 | hypothetical protein | nucleus - chromatin | -3.5839 | -5.7991 | -6.065 | -6.7366 | -3.76 |
| TGGT1_227970 | histone family DNA-binding protein | apicoplast | -3.7168 | -5.2183 | -6.3013 | -5.9505 | -2.15 |
| TGGT1_254800 | hypothetical protein | mitochondrion - soluble | -3.8324 | -5.9067 | -6.627 | -5.7992 | -3.51 |
| TGGT1_218770 | carrier superfamily protein | mitochondrion - membran | -4.0762 | -6.6843 | -5.9854 | -7.1681 | -4.06 |
| TGGT1_271400 | hypothetical protein | N/A | -4.3247 | -6.4516 | -6.922 | -6.3193 | -4.91 |
| TGGT1_233930 | CAF1 family ribonuclease | mitochondrion - soluble | -4.4181 | -6.3053 | -7.4416 | -7.3763 | -4.53 |
| TGGT1_267570 | hypothetical protein | N/A | -4.4512 | -6.2532 | -7.562 | -7.6696 | -4.14 |
| TGGT1_243780 | hypothetical protein | nucleus - chromatin | -4.4763 | -5.7716 | -6.4118 | -6.8919 | -4.68 |
| TGGT1_229620 | hypothetical protein | mitochondrion - soluble | -4.5801 | -6.1847 | -6.6243 | -6.7579 | -4.47 |
| TGGT1_243440 | histone lysine acetyltransferase GCN5-B | nucleus - chromatin | -4.5968 | -6.8496 | -7.2321 | -6.7817 | -2.96 |
| TGGT1_261070 | apicoplast triosephosphate translocator APT1 | apicoplast | -5.0724 | -7.53 | -7.9403 | -7.6782 | -4.44 |
| TGGT1_236210 | peptidase M16 family potein, putative | mitochondrion - membran | -5.1003 | -7.2078 | -7.9214 | -7.4742 | -4.74 |
| TGGT1_256970 | vacuolar ATP synthase subunit A, putative | PM - peripheral 2 | -5.3417 | -6.917 | -8.1861 | -7.6989 | -3.61 |
| TGGT1_205470 | translation elongation factor 2 family protein, putative | cytosol | -6.2901 | -7.8083 | -8.328 | -8.1984 | -4.31 |

To validate whether the genes identified as more fitness conferring in the mutants were true hits, we tested six genes (TGGT1_308830, TGGT1_219190, TGGT1_254840, TGGT1_262640, TGGT1_279390, and TGGT1_249270) from among those identified in all three mutants, particularly those with phenotype scores of >-2.0 in WT parasites indicating knock-out feasibility. WT, *Δgra14*, or *Δcpl* parasites were transfected with sgRNAs targeting the six genes. sgRNAs targeting *sag1* and *cdpk1* were included as negative and positive controls, respectively^31^. Nascent transfected populations were directly plaqued in the presence of solvent (ethanol, no drug selection) or 3 µM pyrimethamine (drug selection) for 7 or 10 days, respectively. The wells were fixed, stained with crystal violet, and imaged. Using a modified Cellpose pipeline, we automated measuring the number and size of plaques (Fig. S3A). The no-drug selection wells served as a correction factor to determine the number of viable parasites within drug selected wells.

As expected, the *Δcdpk1* wells produced few to no plaques since *CDPK1* is an essential gene^31^. This result confirms that disrupting fitness conferring genes reduced the WT and mutants’ fitness. There were trends toward Δ*gra14*Δ*254840* and Δ*cpl*Δ*254840* having lower plaque efficiencies than WTΔ*254840* (Fig. S3B), but the differences were not statistically significant. Compared to Δ*sag1*, WTΔ*262640* plaques were statistically smaller while there was a similar trend in the Δ*gra14*Δ*262640* plaque size. Additionally, Δ*247290* parasites trended towards smaller plaque sizes in both WT and Δ*gra14* parasites. The smaller plaque size in the WT strain for these mutants suggest these genes are false positives (Fig. S3C). Notably, in previous screens the phenotype scores of Δ*262640* and Δ*247290* were lower (-4.0 and -4.8, respectively) than what we observed (-0.8 and -2.2, respectively). None of the other genes had significantly smaller plaque sizes, indicating an inability to conclusively identify synthetic lethal genes. However, we verified that disrupting fitness-conferring genes reduced the WT and mutants’ fitness.

We reexamined our data to assess patterns in the ingestion mutant pairs. The largest overlap in significant genes was between Δ*cpl* and Δ*crt* with 201 total genes, likely due to both proteins being in the PLVAC. Thus, some hits could be involved in compensating for phenotypes associated only with the PLVAC. The Δ*gra14* and Δ*cpl* mutants and the Δ*gra14* and Δ*crt* mutants shared 76 and 36 genes, respectively. This suggests these genes probably do not function together in other pathways outside the ingestion pathway (Fig. 1C).

To determine if this overlap was significant or due to random chance, we tested hypergeometric overlap and found significant overlap in all three gene pairs. This led us to conclude that our CRISPR screens successfully identified genes that were more fitness conferring due to the disruption of the ingestion pathway or the PLVAC.

To identify potential compensatory mechanisms among the more fitness-conferring genes, we conducted a metabolic pathway analysis using tools available on ToxoDB. No clear compensatory pathway emerged from the 30 genes more fitness-conferring shared by all three mutants (Table S2). Thus, we broadened our search to include pathway pairs to determine if a higher number of genes could reveal more fitness-conferring metabolic pathways.

The Δ*cpl* and Δ*crt* samples had the largest overlap of shared hits. Interestingly, all six genes required for pyrimidine biosynthesis (TGGT1_210790, TGGT1_259660, TGGT1_259690, TGGT1_291640, TGGT1_293610, TGGT1_308580) were among the overlapping genes. Also, four overlapping genes are involved in fatty acid (FA) biosynthesis (TGGT1_217740, TGGT1_221320, TGGT1_225990, TGGT1_251930), eight in the TCA cycle (TGGT1_215280, TGGT1_219550, TGGT1_244200, TGGT1_290600, TGGT1_305980, TGGT1_309752, TGGT1_313140, TGGT1_314400), and five in lysine degradation (TGGT1_219550, TGGT1_236570, TGGT1_244200, TGGT1_301120, TGGT1_305980) Fig. 2A, Table S2).

**Figure 2.**
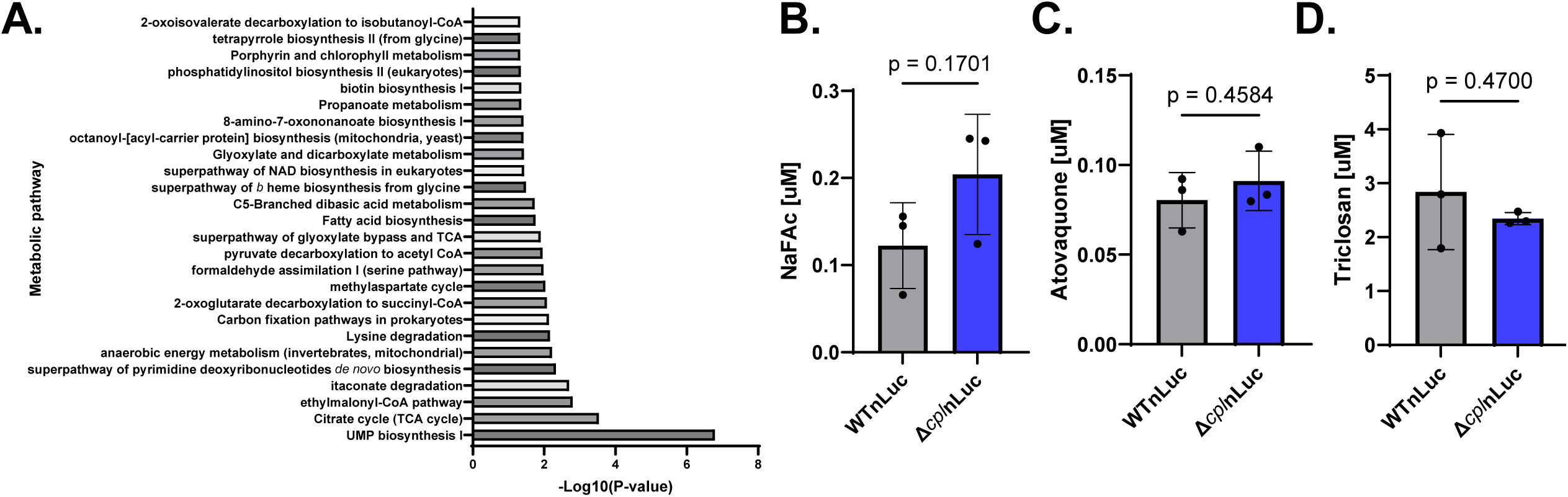
Ingestion mutants rely on several metabolic pathways. **(A)** Pathway enrichment analysis of the significantly more fitness conferring hits shared between the RHCas9Δ*cpl* and RHCas9Δ*crt* screens showing Kegg and Metacyc hits from ToxDB using default settings, with duplicate pathways removed and single gene pathways excluded. **(B, C, D)** Confluent HFF monolayers in 96 well plates were infected in triplicate with 2,000 WTnLuc or Δ*cpl*nLuc parasites in different concentrations of the TCA-cycle inhibitor sodium fluoroacetate (NaFAc) (B), the electron transport chain inhibitor atovaquone (C), or the FA biosynthesis inhibitor triclosan (D). After 4 days at days at 37 °C in 5% CO_2_ parasite growth was measured by luciferase activity. Luminesce values were normalized to growth in vehicle (DMSO) and the IC_50_ was calculated for parasite growth in each drug. The dots are mean IC_50_ values with SD as the error bars from 3 biological replicates. Mann-Whitney test.

The gene pairs between the Δ*gra14* mutant and the Δ*cpl* and Δ*crt* mutants did not show significant impacts on metabolic pathways, with the pathway analysis returning only 1-2 genes per pathway (Table S2). This data suggests that the Δ*cpl* and Δ*crt* mutants might be more reliant on their own biosynthetic pathways due to decreased turnover of material in the PLVAC. In contrast, reduced uptake through the ingestion pathway in Δ*gra14* parasites does not appear to increase the parasites’ reliance on biosynthetic pathways.

To determine whether PLVAC disruption increased reliance on metabolic processes, we tested the ability of RHΔ*ku80*Δ*hxg*nLuc:*hxg* (WTnLuc) and RHΔ*ku80*Δ*hxg*nLuc:*hxg*Δ*cpl:dhfr* (Δ*cpl*nLuc) parasites^32^ to grow in the presence of drugs targeting these pathways. Nano luciferase expression allowed us to measure parasite growth via luminescence. We hypothesized that increased reliance on biosynthetic pathways would heighten sensitivity to their disruption. However, WTnLuc and Δ*cpl*nLuc parasites did not have significantly different IC_50_ values when grown in the presence of sodium acetate, atovaquone, or triclosan, which target the TCA cycle, the electron transport chain (ETC), and FA biosynthesis, respectively (Fig. 2B–D). While Δ*cpl* was not more susceptible to these drugs, it remains unclear if there was an effect on the metabolomic profile of these parasites.

Our metabolic pathway analysis motivated us to conduct bulk metabolomics on the ingestion mutants to determine the direct impact of the ingestion pathway loss on the resources available to the parasite. We harvested WT, Δ*gra14*, Δ*cpl*, and Δ*crt* mutants after 44 hours of growth in confluent HFF monolayers. Additionally, we included two WT samples treated with LHVS, a chemical inhibitor of CPL, and DMSO, as a vehicle control – providing comparative points for our WT and Δ*cpl* samples.

Principal component analysis (PCA) of detected metabolites showed that WT and WT-DMSO clustered together, while Δ*gra14*, Δ*cpl*, and Δ*crt*, and 4 out of 6 WT-LHVS samples clustered together and away from the WT samples, indicating conserved changes in these parasites (Fig. S4A). Due to two outliers in the WT LHVS data, we removed them from the analysis. We calculated the Log2 fold change of all our metabolites in the ingestion mutants to the WT condition and the LHVS to the DMSO condition (Table S3 – Log2FC Meta, Fig. S4B). However, this left us with many metabolites that did not slot into pathways detected in our pathway analysis so we restricted further analysis to the central carbon metabolism, pentose phosphate pathway (PPP), the citric acid cycle (TCA), energy molecules, nucleosides, and amino acids. We noticed that most metabolites in the Δ*cpl* and LHVS samples followed similar trends (Fig. S4C), but were not identical, possibly due to LHVS affecting host CPL and thus impacting the resources available for parasitic scavenging.

Continuing the analysis with only the WT, Δ*gra14*, Δ*cpl*, and Δ*crt* samples, we observed several trends from our bulk metabolomics data. Intermediates and end products of glycolysis were reduced in the ingestion mutants (Fig. 3A–B). Specifically, pyruvate was significantly decreased in all three mutants, and glucose was significantly reduced in Δ*gra14* and Δ*cpl* mutants. Within the PPP 6-O-phosphono-D-gluconic acid was increased in all the ingestion mutants. Since ribose is a main product of the PPP, these findings suggest the need for increased pyrimidine biosynthesis due to disrupted nucleoside salvage. This was further supported by significantly lower levels of guanosine, adenosine, and cytidine in Δ*gra14* parasites (Fig. 3A). Precisely how the ingestion pathway is connected to nucleoside salvage remains to be determined.

**Figure 3.**
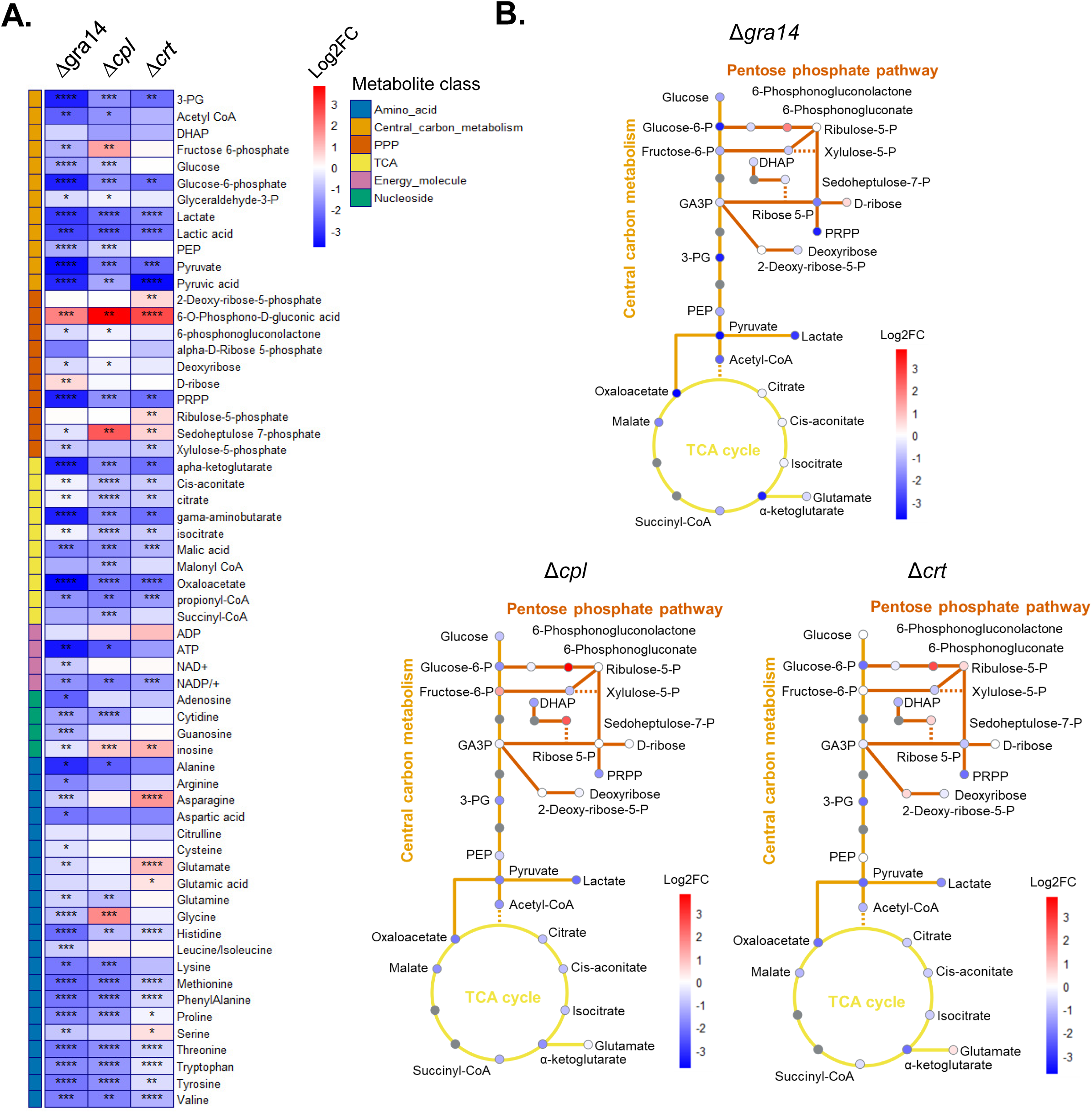
Metabolic analysis of ingestion mutants shows decrease in amino acids and glycolic end products. **(A)** Heat map of select metabolites from bulk metabolomics data for the ingestion mutants, RHCas9Δ*gra14,* RHCas9Δ*cpl,* RHCas9Δ*crt*. Parasites infected HFFs for 44 h before the monolayer was harvested. Parasites were freed from host cells and washed to remove residual media then subjected to LC-MS. Each metabolite was normalized to the RHCas9WT condition then the Log2FC was calculated. * *p*<0.5, ** *p*<0.01, *** *p*<0.001, **** *p*<0.0001, t-test. *n*=6 biological replicates. **(B)** Metabolic pathway schematic for RHCas9Δ*gra14,* RHCas9Δ*cpl, and* RHCas9Δ*crt* depicting the central carbon metabolism (glycolysis and fermentation) in orange feeding into the citric acid cycle in yellow or the pentose phosphate pathway in dark orange. The color of the dot is the Log2FC that metabolite experienced in the mutant parasite. Grey dots are metabolites that were not looked at in our analysis.

The ingestion mutants exhibited markedly reduced levels of most amino acids, with Δ*gra14* mutants showing significantly reduced levels of all amino acids except alanine and valine, which still trended lower (Fig. 3A). The Δ*cpl* mutant had higher levels of glycine, while leucine and isoleucine trended upward. While the Δ*crt* mutant showed higher levels of glutamic acid, serine, and three other amino acids trending upward. This data supports the role of the ingestion pathway in scavenging host cytosolic proteins. Despite fewer amino acids in the mutants, there is little to no growth defect compared to WT parasites, suggesting the parasite has an excess pool of amino acids and that it can acquire sufficient amino acids through other means such as amino acid transporters.

To assess if the ingestion pathway significantly contributes to amino acid acquisition under nutrient limiting conditions, we grew the mutants in single amino acid-depleted media. Using the nano-luciferase strain, we generated Δ*gra14*nLuc and Δ*crt*nLuc strains by disrupting the gene locus and replacing the coding sequence with a *dhfr* resistance cassette, like what was done for the Δ*cpl*nLuc strain^32^ (Fig. S5). Starving HFFs for 24 hours before parasite infection ensured reduced internal amino acid stores. We focused on tryptophan and phenylalanine, which were significantly reduced in all three mutants, and included tyrosine and arginine due to the tyrosine being the other aromatic amino acid, host arginine levels being controlled by host cells, and *Toxoplasma* being auxotrophic for both amino acids^2^.

We assayed parasite growth in complete (D1) and tryptophan-free (D1 -W) media to determine doubling time over 3 days. There was no difference in doubling time for WTnLuc, Δ*gra14*nLuc, Δ*cpl*nLuc, and Δ*crt*nLuc parasites in complete media. However, in tryptophan-free media, Δ*cpl*nLuc and Δ*crt*nLuc mutants grew significantly slower than WTnLuc parasites (Fig. 4A). Parasites did not grow in phenylalanine, arginine, or tyrosine-free media, so we tested diminishing concentrations of these amino acids on WTnLuc parasites to identify minimal growth conditions (Fig. S6). Testing parasites in 6.5 μM F, 8.6 μM R, or 3.4 μM Y showed smaller shifts in overall doubling time, yet Δ*cpl*nLuc still grew significantly slower in arginine- and tyrosine-limiting conditions (Fig. 4B). Additionally, all nutrient limited growth conditions grew slower than the complete media (Fig. 4A–B). Overall, this data suggests that *Toxoplasma* may rely on material turnover in the PLVAC during nutrient-limited situations, facing difficulties in resource scavenging.

**Figure 4.**
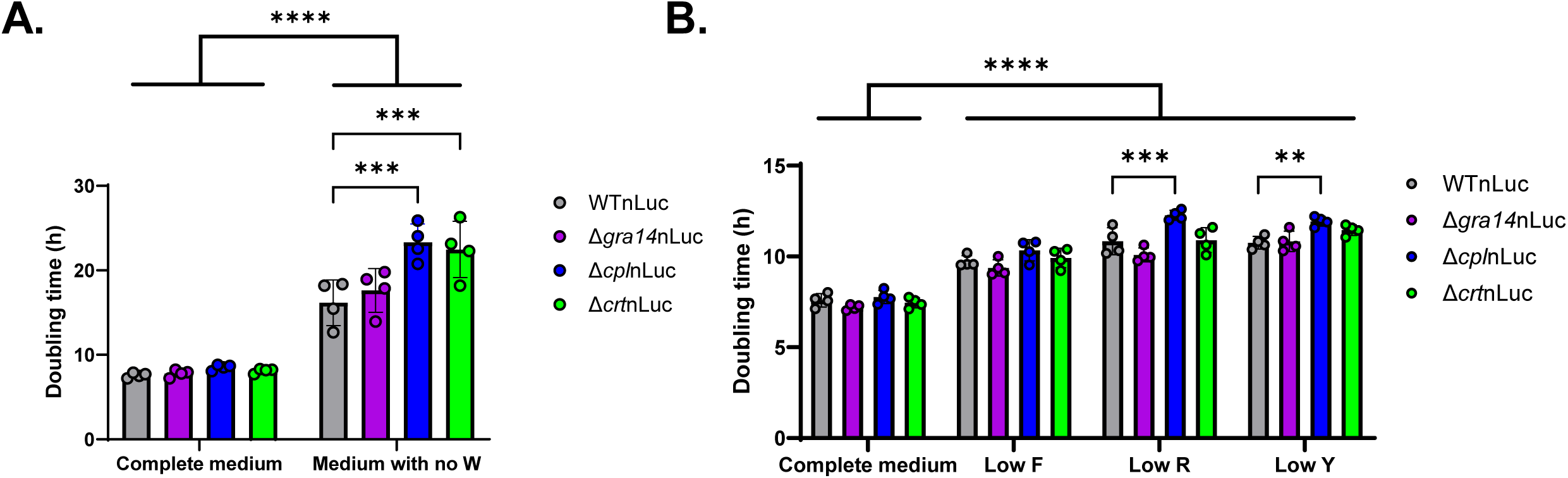
Nutrient limiting conditions inhibit ingestion mutant replication. **(A, B)** HFF monolayers grew to confluency over 1 week in 96-well plates. 24 h prior to infection D10 was replaced with complete D1 or D1 without tryptophan (A) or with 6.5 µM phenylalanine, 8.6 µM arginine, or 3.4 µM tyrosine (B). 2,000 WTnLuc, Δ*gra14*nLuc, Δ*cpl*nLuc, or Δ*crt*nLuc parasites infected monolayers in triplicate per condition. Luciferin was added at 4, 24, 48, and 72 hours post infection to measure parasite growth. Samples were normalized to the 4hpi reading then Log2-transformed. The slope of the linear regression is the doubling time of the parasites. The dots represent mean doubling times with SD as the error bars. ** *p*<0.01, *** *p*<0.001, **** *p*<0.0001 two-way ANOVA with Tukey’s multiple comparison. *n*=4 biological replicates.

## Discussion

Our results provide insights into the adaptive mechanisms employed by *Toxoplasma* in the absence of a functional ingestion pathway. Despite potential bottlenecks in our CRISPR screens, comparison of the WT screen done herein to the original screen by the Lourido Lab^27^ showed a strong correlation. Further, we noted statistically significant overlaps in hits shared by pairwise analysis of Δ*gra14*, Δ*cpl*, and Δ*crt*. The overlap between Δ*cpl* and Δ*crt* was particularly strong and involved pathways for pyrimidine biosynthesis, FA biosynthesis, the TCA cycle, and lysine degradation. This suggests a pivotal compensatory role for these biosynthetic pathways when the ingestion pathway is disrupted. However, we were unable to confirm increased reliance on the pathways identified via metabolic pathway analysis, via our drug sensitivity assay. Our bulk metabolomic profiles of the ingestion mutants complemented our pathway analysis findings, revealing a pronounced deficiency in glycolytic intermediates and certain amino acids, suggesting a marked alteration of these biosynthetic pathways when the ingestion pathway is compromised.

Our detailed metabolomic analysis supports the hypothesis that loss of the ingestion pathway significantly impacts amino acid acquisition. Mutant strains exhibited a global decrease in most amino acids, indicative of disrupted amino acid salvage from the host cytosol. Notably, the Δ*gra14* mutant displayed markedly reduced levels of all amino acids except alanine and valine, while the Δ*cpl* and Δ*crt* mutants showed a slightly more complex but similar amino acid profile disruption. This supports the notion that the ingestion pathway might function in scavenging essential nutrients, particularly amino acids, from the host.

We explored this hypothesis by growing the mutants in single amino acid-depleted media. The results showed that, under nutrient limitation, PLVAC mutants displayed growth impairments, especially in conditions deficient in tryptophan, arginine, and tyrosine. This confirms that *Toxoplasma* uses the PLVAC as a site of protein turnover for amino acids during host nutrient deprivation, such as during host infection-induced amino acid starvation.

Our study provides valuable insights into the adaptive resilience and metabolic flexibility of *Toxoplasma*. This is highlighted in the metabolomic data and pathway analysis that showed a decrease in nucleosides and an increased reliance on pyrimidine biosynthesis in these mutants. This data suggests that the ingestion pathway may also scavenge host cytosolic nucleosides, possibly in the form of host rRNA, mRNA, and tRNA. Recently a preprint study localized TgENT1, an essential equilibrative nucleoside transporter, to the PLVAC^33^. This transporter could play a role in liberating host nucleosides that were acquired by the parasite via the ingestion pathway.

Among the limitations of this study, we were unable to directly validate any of the overlapping hits as being synthetically lethal. Of the 30 genes that were consistently more fitness conferring in all three ingestion-deficient mutants, only 9 of the hits had WT phenotype scores that were indicative that they might be dispensable in WT parasites. However, upon looking at data from previous screens we can see that 5 of these genes, TGGT1_ 262640, TGGT1_221320, TGGT1_286530, TGGT1_288880, and TGGT1_249270 have phenotype scores <-2, which indicates another screen found them fitness conferring in WT parasites^27^. This highlights the importance of recognizing a degree of uncertainty with phenotype scores. Also, we attempted to validate the hits by measuring viability (plaquing efficiency) and overall progression through the lytic cycle (plaque size). These assays do not measure fitness in the same way as the CRISPR screen, which is essentially a competition assay. Alternative ways of validating hits for synthetic lethal screens including by pairwise competition assays should be considered.

Our findings underscore the reliance of *Toxoplasma* on a multiplicity of adaptive pathways to compensate for the loss of nutrients. This flexibility ensures that the parasite can thrive under various constraints, highlighting its evolutionary success as an intracellular pathogen.

## Materials and Methods

### Host cells and cell culture

Human foreskin fibroblasts, HFFs, (ATCC) were maintained in Dulbecco’s modified Eagle’s medium (DMEM) with 4.5 g/L glucose and supplemented with 2mM L-glutamine, 10% Cosmic calf serum, 20 mM HEPES, 5 µg/mL pen/strep (D10) at 37°C and 5% CO_2_ unless otherwise stated. *Toxoplasma gondii* strains were passaged in the above HFFs.

### Parasite strain generation

A list of all parasite strains used in this study can be found in Table S4 – Strains, a list of all primers can be found in Table S5 – Primers, and a list of all sgRNAs can be found in Table S6 – Plasmids. To generate RHCas9*Δgra14*, RHCas9 parasites were transfected with 50 µg of circular pU6-DHFR-GRA14gRNA1 which targets the first exon of the gene. To prevent integration of the resistance maker the plasmid was not linearized. Media was changed 24 h later. Once the HFF monolayer was lysed parasites were syringed and filtered before being diluted into 96- well plates for single clones. Parasites were cloned for one week before being expanded into 24-well plates to allow for sufficient material generation for DNA extraction. DNA was extracted using Qiagen DNeasy Blood and Tissue kit. PCR was performed using Q5 polymerase with primers, P1 and P2, that flanked the cut site targeted by the gRNA. PCR fragments were purified using Qiagen PCR clean up kit and were sent for Sanger sequencing to identify INDEL mutations that would knock the genes out of frame. This same procedure was used for generating RHCas9Δ*crt* using pU6-DHFR-CRTgRNA 1 and primers 3 and 4. Due to difficulties acquiring RHCas9Δ*cpl* a similar procedure was used but a plasmid was chosen that would allow for enrichment of transfected parasites, pCas9-GFP-Ble-CPLgRNA1. Twenty-four hours after transfection the monolayer was scraped, syringed, and filtered. Parasites were resuspended in PBS and 5%FBS and the population was flow sorted and the GFP expressing parasites were kept and infected a monolayer. Once this monolayer was lysed the procedure above was followed and primers 5 and 6 were used to verify the INDEL. Protein disruption was validated by western blotting using antibodies specific to these proteins.

To generate RHΔ*ku80nLuc*Δ*gra14*, RHΔ*ku80*nLuc parasites were transfected with 50 µg of pCas9-GFP-Ble-GRA14gRNA1 and 10 µg of repair template that had 40 bp of homology to the 5′ and 3′ of the genomic DNA, flanking a DHFR resistance cassette. Twenty-four hours post transfection media was changed and 3 µM pyrimethamine as added for selection. Once the population was lysing, the monolayer was scraped, syringed, and filtered then diluted into 96 well plates to generated single clones. After one week, the Phire Tissue Direct PCR kit was used to test for integration of the DHFR resistance cassette using primers P9 and P10 for 5′ integration. This was also validated at the protein level by western blotting. A similar approach was used to generate RHΔ*ku80nLuc*Δ*crt*, however two sgRNAs, pCas9-GFP-Ble-CRTgRNA1 and pCas9-GFP-Ble-CRTgRNA2, were used as *CRT* is a multi-exon gene so sgRNAs targeting the N and C terminus of the gene would increase the efficiency of the knockout process. Primers P3 and P10 were used to test for 5′ integration.

### Ingestion assay

Inducible mCherry HeLa cells and harvesting steps were used from a previous study^17^. After harvesting parasites were fixed with 4% methanol-free formaldehyde prior to imaging, using a Zeiss Axiovert Observer fluorescence microscope. Samples were blinded and mCherry positive parasites were counted. Data was imported to PRISM for graphing and statical analysis.

### Immunoblot

A list of all antibodies used in this study can be found in Table S7 – antibodies. Parasite lysates were harvested using RIPA buffer solution (Thermo Scientific, 89900) treated with EDTA-free miniComplete protease inhibitor (Roche, 11836153001). PVDF membranes were activated with methanol prior to transferring proteins onto the membrane. Membranes were blocked with 5% milk in PBS-tween prior to primary antibody incubation O/N at 4 °C, or at 1 h at RT for loading controls, and then incubated with horseradish peroxidase (HRP) conjugated secondary antibodies for 1 h at RT. For detection, the membranes were incubated for 2 min with the SuperSignal West Pico PLUS Chemiluminescent substrate or SuperSignal West Femto Maximum sensitivity substrate. Chemiluminescent signal was visualized using the Syngene PXi6 imaging system. Membranes were stripped using 4% trichloroacetic acid after which they would be blocked and probed again.

### CRISPR screens

The genome wide screens were performed using an sgRNA library (CrLib) containing 10 sgRNAs per gene for 8,156 genes in *Toxoplasma*. We followed the established protocol^26^ with few changes. The CrLib was linearized using AseI and split into 12 cuvettes for transfections of 50 µg of linearized plasmid with 5×10^7^ parasites each. Twelve D150 dishes of HFFs were infected with the parasites and 24 h post infection, media was changed to D10 with 40 µM chloramphenicol, 3 µM pyrimethamine, and 10 ug/mL DNase1. After the final passage where 1×10^8^ parasites were pelleted, and genomic DNA was extracted from the parasites using the Qiagen DNeasy Blood and Tissue kit. To prepare the samples for sequencing sgRNAs were amplified from the genomic DNA using the primer pairs listed in Table S4 – Primers, gel purified and sequenced on an Illumina Nova Seq. Three screens for each mutant and the WT were done sequentially.

### Data analysis

A custom KNIME workflow was used to trim the sequencing reads so they began with the sgRNA sequence, and were compatible with the Countess R script (https://github.com/LouridoLab/CRISPR_Analysis), detailed in the protocol paper^26^. The phenotype score is then calculated as the log2 fold change for the 5 most abundant gRNAs for each gene. To determine which genes underwent a change in phenotype score in the mutants the average phenotype score in the WT and mutants were compared and an average phenotype shift was determined. This shift was determined to be significant if it was greater than 2 times the standard deviation the average shift in the 497 sexual genes^30^ experienced in the mutant compared to the WT. Additionally, one-sided Wilcoxon signed rank test was used to determine the probability of each gene having a significant shift compared to the shift in the 497 sexual genes, FDR<0.01. Genes that were determined as significant if both methods were classified as hits. To determine if there was statistically significant overlap between screens, we used hypergeometric overlap testing (http://nemates.org/MA/progs/overlap_stats.html).

### Gene enrichment analysis

Genes that were identified as significantly shifted were subject to gene enrichment analysis using the metabolic pathway enrichment analysis from ToxoDB, using default settings. These results can be found in Table S2.

### Validation of screens

Thirty genes were fitness conferring genes in all three mutants. Of these 6 genes had fitness scores that were above -2.5 in the WT suggesting they were dispensable. To validate if these hits were truly synthetic lethal hits the genes would be disrupted in the RHCas9WT, RHCas9Δ*gra14*, and RHCas9Δ*cpl* parasites and fitness of the resulting single or double mutants would be determined. For a negative control, the pU6-DHFR-SAG1 sgRNA was used. For a positive control an sgRNA for CDPK1 was designed to test for lethality. The pU6-DHFR back bone (from above) was used and the sgRNAs were designed to target the first exon of each gene. Like above 5×10^7^ parasites were transfected with 50 µg of linearized plasmid. Immediately after transfection parasites were diluted in D10 and plated on 6-well plates, in duplicate. Wells that were treated with ethanol were seeded with 100 and 1,000 tachyzoites, wells treated with 1.5 µM pyrimethamine were seeded with 1,000 or 10,000 tachyzoites. Ethanol-treated plates grew for 8 days while pyrimethamine-treated plates grew for 10 days. On the final day of growth, supernatant from the pyrimethamine-selected RHCas9WT plates was collected for validation of the gene disruption (detailed below). Then monolayers were washed with PBS, fixed with 4% methanol free formaldehyde, and stained with 2% crystal violet in 20% ethanol. Wells were imaged on a Nikon Ti-2E microscope at 4x magnification before images were stitched together using NIS Elements and exported as TIFFs for analysis. Two wells were analyzed per condition. Plaques were detected and quantified using the plaque assay module in spacr (https://github.com/EinarOlafsson/spacr, https://pypi.org/project/spacr/, v. 0.3.62). First, plaques were manually annotated using the spacr module spacr.make_masks in 100 whole well stitched images acquired with the Nikon Ti-2E. The cellpose cyto model^34^ was fine-tuned^35^ on 90 images with the spacr.train module (Table S9 – spacr settings). 10 images were reserved for testing. Once the plaque counts and sizes were computed by spacr, the data was imported to R where in outlier analysis was conducted and the average plaque size and count for each condition was calculated, normalized to the EtOH and Δ*sag1* controls. The data was exported to PRISM for graphing and statistical analysis.

To determine how successful this knockout strategy was, the supernatant of parasites that created plaques was collected and plated for single clones in 96-well plates. After 7 days the Phire Tissue Direct PCR kit was used to probe for disruption of the gene through the integration of the sgRNA fragment at the targeted gene locus with genomic and sgRNA primer pairs (P57–P70, and P55 and P56). Integration of the sgRNA could be detected by running the genomic-sgRNA fragment reactions on a gel. For clones that did not produce a genomic-sgRNA PCR fragment, genomic primer pairs that flanked the cut site were used. That PCR fragment was cleaned up using the Qiagen PCR clean up kit and sent for Sanger sequencing to identify clones with an INDEL.

### Luciferase growth assay

To determine if Δ*cpl*nLuc parasites were more susceptible to drugs that targeted the metabolic pathways that were more fitness conferring in the Δ*cpl* and Δ*crt* mutants, we employed a growth assay. Briefly, 96-well plates were seeded 1 week prior to infection and were allowed to grow to confluency. On the day of infection WTnLuc, and Δ*cpl*nLuc parasites were harvested from monolayers by scraping, syringing, filtering, and centrifugation at 1,200 g for 10 min. The pellet was washed once in HBSS and resuspended again in HBSS. Triplicate wells for each strain for drug concentration were infected at 2,000 tachyzoites per well. Parasites invaded for 4 h at 37 °C in 5% CO_2_ before the wells were washed once with PBS and fresh media was added which contained either DMSO control or a 3-step dilution of sodium fluoroacetate starting at a concentration of 5.4 µM, atovaquone starting at a concentration of 5.4 µM, or triclosan starting at a concentration of 17 µM. Samples were grown for 4 days at 37°C in 5% CO_2_. To assess parasite growth on day 4 media was aspirated from the wells and monolayers were lysed for 10 min in equal 50% 1×PBS 50% 2×Lysis buffer (100 mM MES, 1 mM CDTA, 0.5% Tergitol, 0.05% Antifoam 204, 150 mM KCl, 1 mM DTT, 35 mM Thiourea) with 12.5 µM h-coelenterazine (NanoLight Technology, 301-10), luciferase substrate. Plates were read on Syngene plate reader using the following settings: Integration time: 1s, Optics: top, Gain: 135, Delay: 100 ms, read height: 4.5 mm. Data was analyzed in PRISM and luminescence values were normalized to the DMSO condition before growth curves were plotted and the IC_50_ was calculated.

To determine if there was a defect in growth in nutrient-limited conditions, we adapted a luciferase growth assay similar to^32^. Briefly, 96-well plates were seeded 1 week prior to infection and were allowed to grow to confluency in D10. Twenty-four h prior to infection the D10 was replaced with amino acid free DMEM (US Biological, D9800-13) supplemented with 1% dialyzed fetal bovine serum, 20 mM HEPES, 5 µg/mL pen/strep and further supplemented as described in Table S8 – Media to make D1 complete, D1 - tryptophan, D1 -phenylalanine, D1 -arginine and D1 -tyrosine. The media was brought to pH 7.4–7.6 before being filter sterilized. If necessary, D1 complete was mixed with an amino acid free media to make a certain concertation. To determine inhibitory concentrations for phenylamine, arginine, and tyrosine triplicate wells of a 2-step serial dilution were infected at 2,000 WTnLuc tachyzoites per well. Parasites invaded for 4 h at 37 °C in 5% CO_2_ before the wells were washed once with PBS and fresh media was added. The first reading of sample growth was gathered at this time. Reading of parasite growth was accomplished as above. For inhibitory concentrations of F, R, and Y samples were assayed after 4 days of growth and normalized to the 4 h post infection reading and plotted as a percentage of the D1 complete growth. A nonlinear curve was fit to the graphs in PRISM to estimate the EC_90_.

For growth assays in nutrient-limited media. WT*nLuc*, Δ*gra14nLuc*, Δ*cplnLuc*, and Δ*crtnLuc* parasites were harvested as above. Triplicate wells for each strain for each time point captured were infected at 2,000 tachyzoites per well in plates containing D1 complete, or D1 with a limited concentration of our experimental amino acid. Readings were taken at 4, 24, 48, and 72 h post infection and were normalized to the 4 h post infection readings for each sample before fold change was calculated. The log2 fold change values were determined and used for linear regression to determine the parasite doubling time in PRISM.

### Metabolomics

HFF monolayers were infected with equal number of RHCas9, RHCas9Δ*gra14*, or RHCas9Δ*cpl*, and RHCas9Δ*crt* parasites. After 2 h of invasion the monolayer was washed with DMEM and infection proceeded for 18 h in D1. Samples were either treated with 1 µm LHVS for 24 h or DMSO as the vehicle control before parasites were syringe lysed and harvested for metabolomic analysis using LC-MS/MS. Untargeted metabolomics was conducted as previously published^36^. Data was analyzed in R to identify outliers then to calculate the log2 fold change and significance of the metabolites between the WT, and ingestion mutants and the DMSO and LHVS conditions.

## Acknowledgments

We thank Tyler Smith for help in troubleshooting our initial library prep. We also thank Christopher Giuliano and Yifan Wang for their insights on CRISPR screen data analysis. We thank Dr. Zhicheng Dou from Clemson University for gifting us the RHΔ*ku80nLuc:HXG* and RHΔ*ku80nLuc:HXG*Δ*cpl:DHFR* strains. We appreciate our colleagues Mary O’Riordan, Joel Swanson, Yoshifumi Nishikawa, David Sibley, and Manlio di Cristina for supplying key reagents. We thank the Advanced Genomics Core at the University of Michigan for sequencing our libraries. We are grateful to the Mass Spectrometry and Proteomics Core Facility at the University of Nebraska Medical Center for performing the metabolomics analysis and providing valuable suggestions. Chat-GPT was used to condense the initial version of this manuscript and BioRender was used to create some figures. This work was supported by grants from the US National Institute of Health, including P20 GM130447-01A1 (L.A.), R01 AI144369 (S.L.), and R01 AI120607 (V.B.C.), and a Boehringer Ingelheim Fonds PhD fellowship to B.M.M.

---

Table S1. Screen data: phenotype score, significance, and gene information from ToxoDB for each gene in the CrLib.

Table S2. Pathway enrichment analysis data from ToxoDB for significantly fitness conferring hits from CRISPR screen shared between Δ*gra14*, Δ*cpl*, Δ*crt* and Δ*cpl*, Δ*crt* and Δ*gra14*, Δ*cpl* and Δ*gra14*, Δ*crt*.

Table S3. Metabolites raw: raw metabolic count for the metabolites detected in the analysis. Log2FC meta: Log2FC and p-values of each metabolite with outliers removed compared to WT for Δ*gra14*, Δ*cpl*, Δ*crt* and DMSO for LHVS samples.

Table S4. Strains: strains used in this study

Table S5: Primers: primers used in this study

Table S6: Plasmids: plasmids used in this study

Table S7: Antibodies: antibodies used in this study

Table S8: Media: media recipes for amino acid supplementation

Table S9: spacr settings: settings used during spacr training and data analysis.

**Figure S1.**
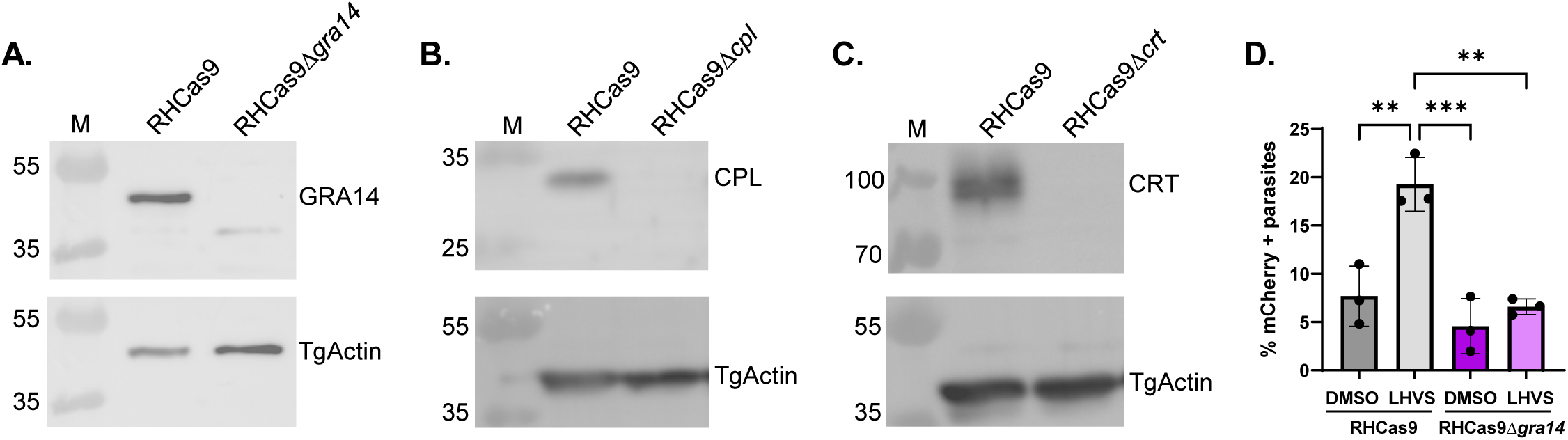
Validation of RHCas9 knockouts. **(A)** Parasite lysates of RHCas9 and RHCas9Δ*gra14* were probed with RbαGRA14 or MsαTgActin (as a loading control). **(B)** Parasite lysates of RHCas9WT and RHCas9Δ*cpl* were probed with MsαCPL or RbαTgActin. **(C)** Parasite lysates of RHCas9WT and RHCas9Δ*crt* were probed with RbαCRT or MsαTgActin. Molecular weight markers (M) are shown in kDa. **(D)** Quantification of host mCherry ingestion 24 h post-infection by RHCas9 or RHCas9Δ*gra14* parasites treated with DMSO (vehicle) or 1 µM LHVS to inhibit degradation of host mCherry in the PLVA(C) ** *p*<0.01, *** *p*<0.001. One-way ANOVA with Tukey’s multiple comparison. *n*=3 biological replicates.

**Figure S2.**
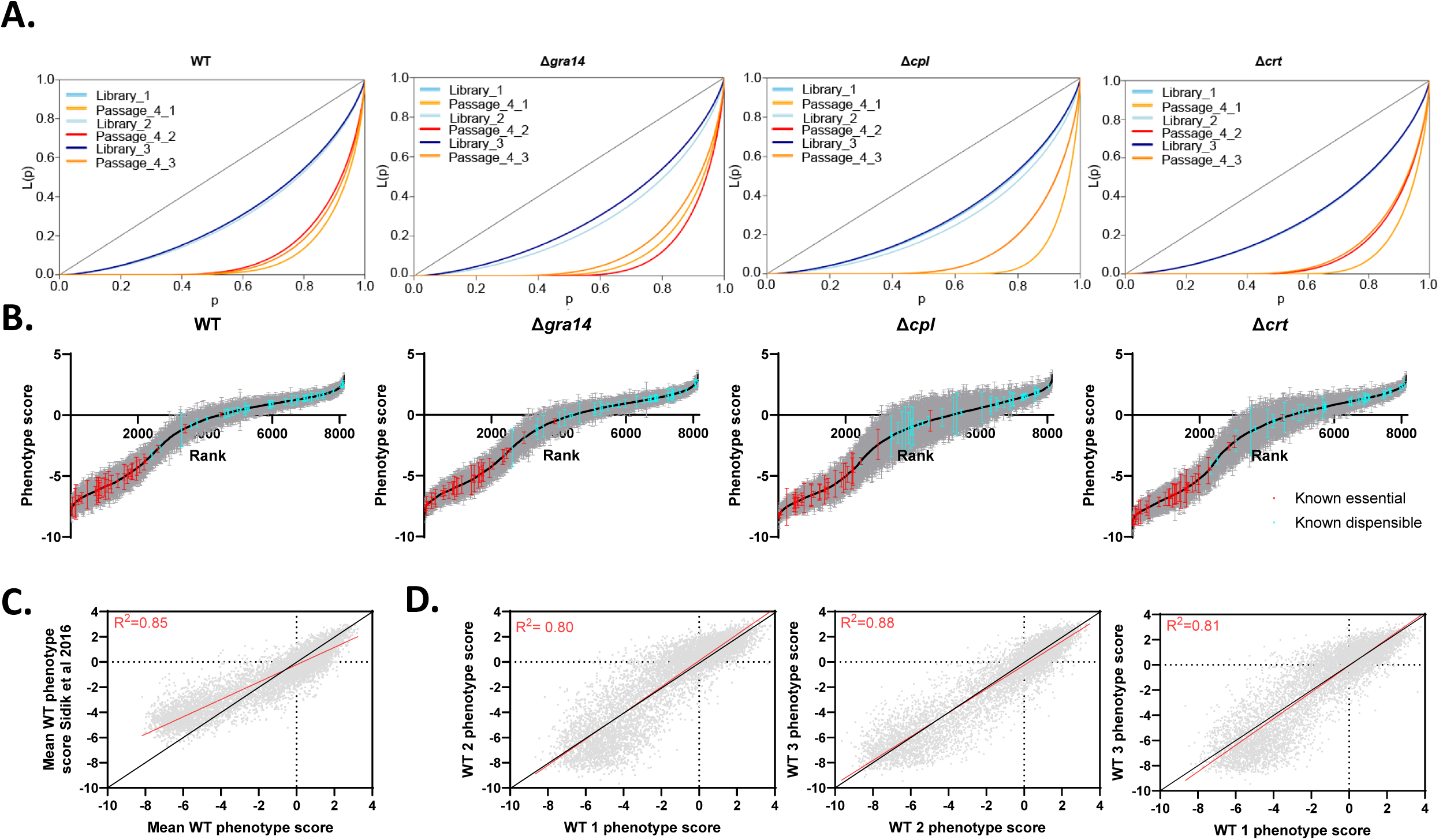
Quality control analysis of genome-wide CRISPR screens. **(A)** Lorenz curves of each CRISPR screen. L indicates the initial library for transfection and P4 is the final passage population. **(B)** Ranked phenotype score graphs with SEM for each screen with known essential (red) and dispensable (cyan) genes marked. **(C)** Correlation of our RHCas9 phenotype scores mapped to the previously published scores from Sidiki et al., 2016. **(D)** Internal correlation of the phenotype score for each WT-screen replicate against the others.

**Figure S3.**
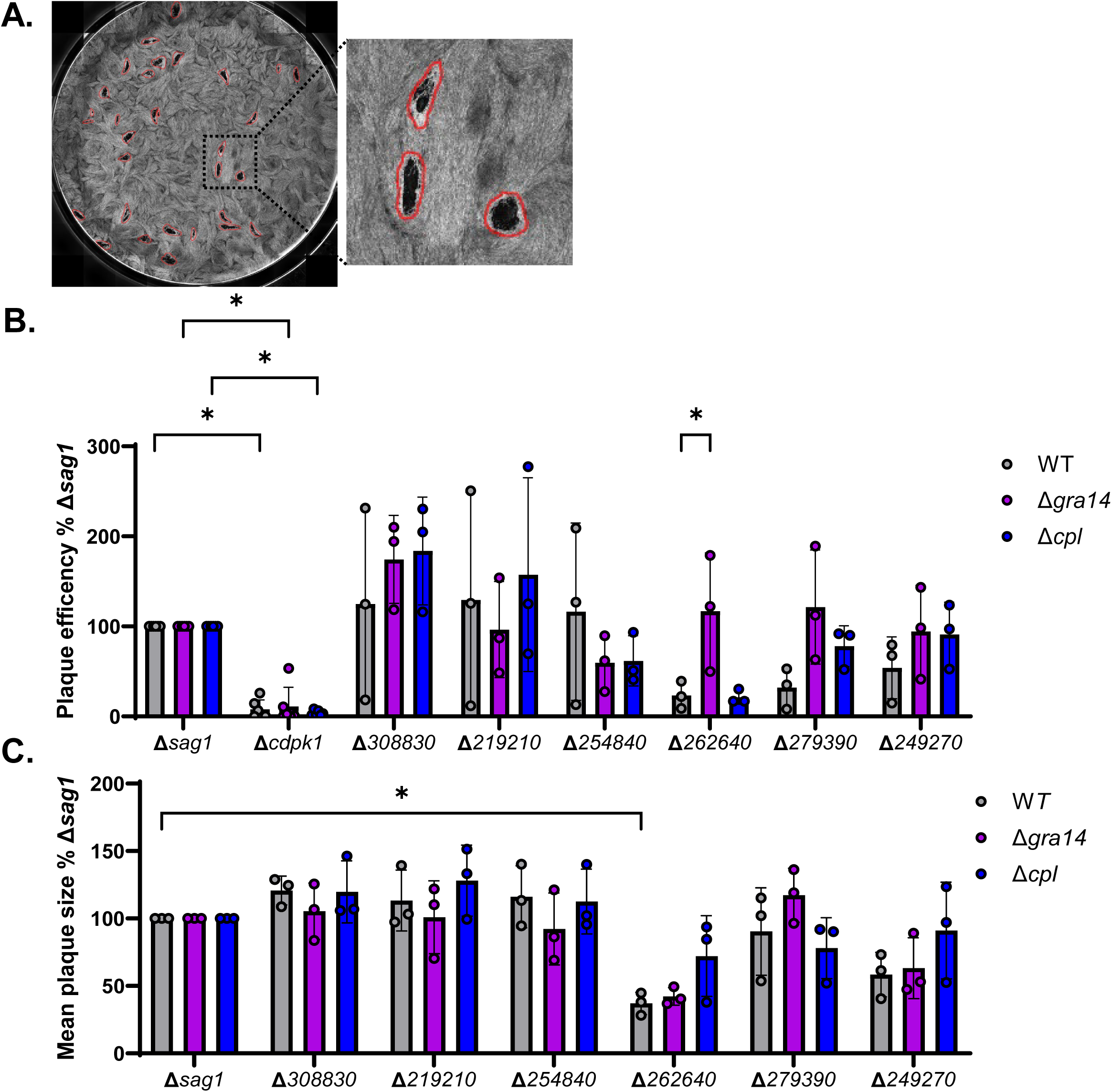
Analysis of plaque efficiency and size for screen hits. **(A)** Representative image of crystal violet stained well after 4x magnification images were stitched together using Nikon Elements. Masks were created after automated plaque identification was performed using the plaque assay module in spacr, a modified cellpose cyto model trained to detect plaques in monolayers of cells, on two wells per condition. Masks were counted for plaque count and size was recorded for plaque size analysis. Red outlines trace mask area onto the input image. **(B)** RHCas9WT, RHCas9Δ*gra14,* RHCas9Δ*cpl* parasites were transfected with an sgRNA targeting the indicated gene, and 1×10^3^ or 1×10^4^ parasites were inoculated into a 6-well plate under 3 µM pyrimethamine selection for 10 days. Wells were fixed in 4% formaldehyde and stained with 2% crystal violet before being imaged. Plaque efficiency was calculated as the number of plaques counted versus the number of parasites added to the well, as determined from the plaque counts of non-drug selected wells, then normalized to the Δ*sag1* value for the given strain. **(C)** Plaque size was calculated and normalized to the Δ*sag1* value for the given strain. **B, C** The dots are means with SD as the error bars. * *p*<0.5, two-way ANOVA with Tukey’s multiple comparison, only comparisons between same strain Δ*sag1*:ΔGOI and WT:ingestion mutant ΔGOI are being shown. n=3, except for Δ*sag1* and Δ*cdpk1* in **B** which was *n*=6

**Figure S4:**
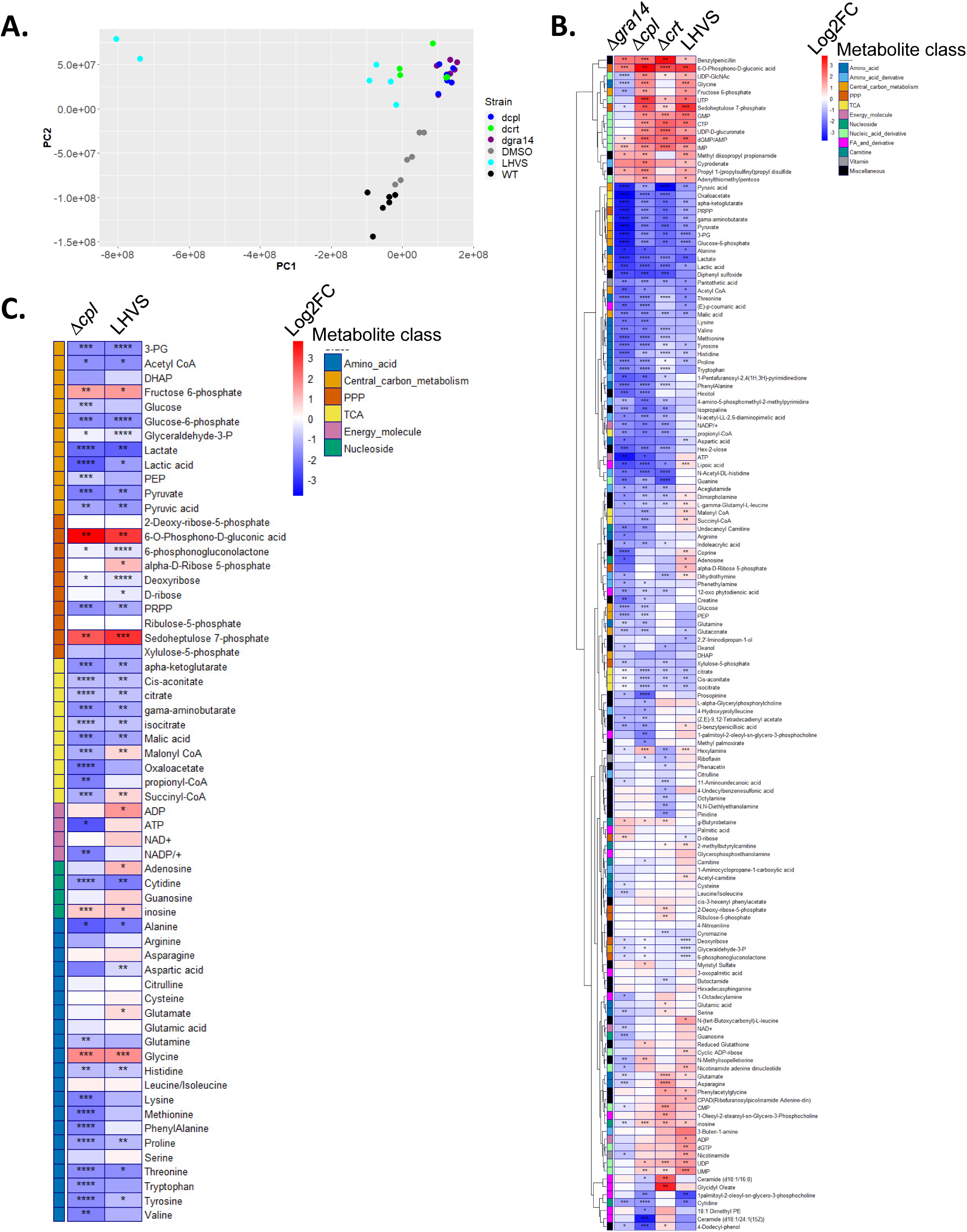
Consistent clustering of groups in metabolic data. **(A)** PCA plot from the RHCas9, RHCas9Δ*gra14,* RHCas9Δ*cpl,* RHCas9Δ*crt*, RHCas9 DMSO and RHCas9 LHVS metabolomics data. *n*=6 **(B)** Heat map of hierarchical clustering of metabolite Log2FC for the ingestion mutants and RHCas9 LHVS normalized to RHCas9 and RHCas9 DMSO, respectively. Parasites infected HFFs for 44 hours, with 24 h of DMSO or LHVS treatment as required, before harvest. Parasites were freed from host cells and washed to remove residual media then subjected to LC-MS. **(C)** Heat map of select metabolites from bulk metabolomics data for RHCas9Δ*cpl,* RHCas9 LHVS. **B, C** The LHVS samples 1 and 4 were removed due to being outliers. Statistical significance was determined by t-test, * *p*<0.5, ** *p*<0.01, *** *p*<0.001, **** *p*<0.0001. RHCas9Δ*gra14,* RHCas9Δ*cpl,* RHCas9Δ*crt n*=6 and RHCas9 LHVS *n*=4.

**Figure S5.**
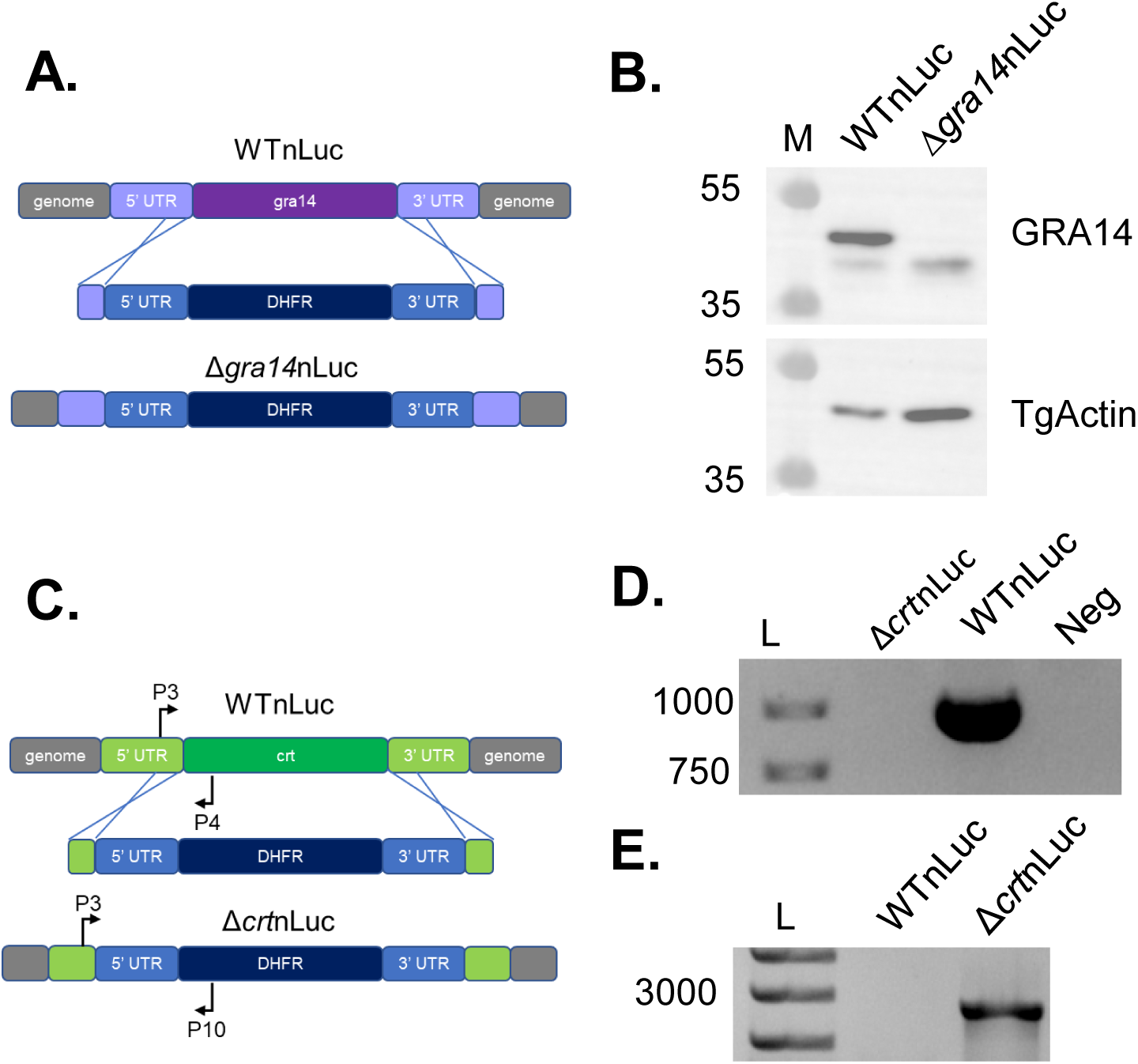
Generation of RHΔ*ku80*Δ*gra14*nLuc. **(A)** Schematic of replacing the *gra14* locus with DHFR **(B)** Parasite lysates of RHΔ*ku80*nLuc, RHΔ*ku80*Δ*gra14*nLuc with RbαGRA14 or MsαTgActin as a loading control. Molecular weight markers (M) are shown in kDa. **(C)** Schematic of replacing the *CRT* locus with DHFR **(D)** Genomic DNA of WTnLuc and Δ*crt*nLuc PCR’d with P3 and P4 to look for the native *CRT* locus. **(E)** Genomic DNA of WTnLuc and Δ*crt*nLuc PCR’d with P3 and P10 to assay 5’ integration. **D, E** DNA Ladder (L) is shown in bp.

**Figure S6.**
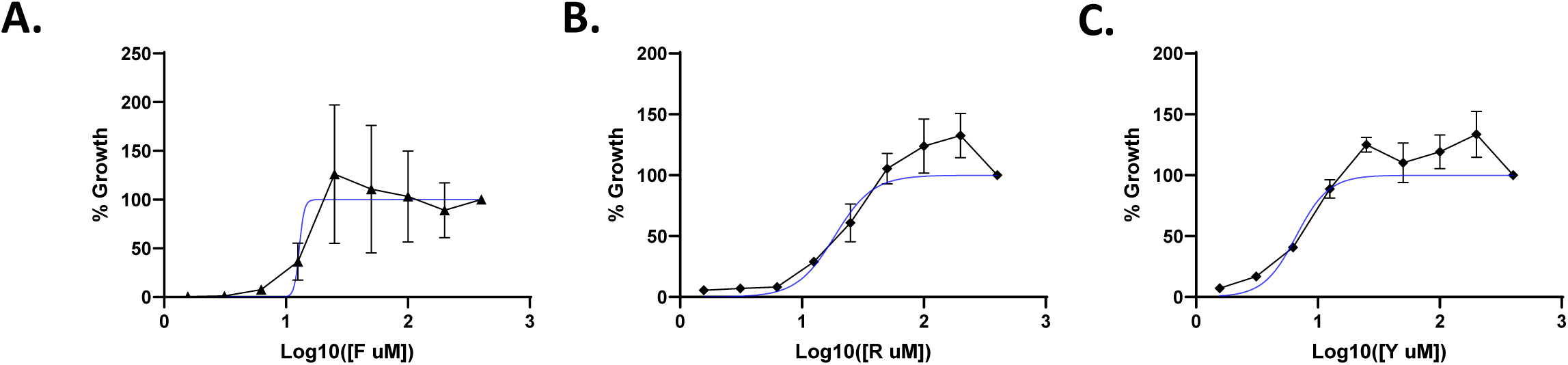
Growth of *Toxoplasma* in phenylalanine-, arginine-, or tyrosine-limiting conditions. **(A, B, C)** Confluent HFF monolayers in 96-well plates were prestarved for 24 hours in D1 complete media serially diluted in D1 missing phenylalanine (A), arginine (B), or tyrosine (C) to create D1 with concertation gradients for the experimental amino acid before infecting with 2,000 WTnLuc parasites per well with 3 technical replicates per dilution. Four hours post invasion, the media was changed keeping the dilution series, and the first reading was taken. The rest of the samples grew for 4 days at 37 °C in 5% CO_2_. Readings were normalized with the 4 hours post-infection infection readings and plotted as a percentage of growth as the complete media. A non-linear fit was used to determine the EC_50_. The dots are the mean growth with SD represented by the error bars. *n*=3 biological replicates.

